# Substrate recognition determinants of human eIF2α phosphatases

**DOI:** 10.1101/2021.07.12.452048

**Authors:** George Hodgson, Antonina Andreeva, Anne Bertolotti

## Abstract

Phosphorylation of the translation initiation factor eIF2α is a rapid and vital cellular defence against many forms of stress. In mammals, the levels of eIF2α phosphorylation are set through the antagonistic action of four protein kinases and two heterodimeric protein phosphatases. The phosphatases are composed of the catalytic subunit PP1 and one of two related non-catalytic subunits, PPP1R15A or PPP1R15B (R15A or R15B). Attempts at reconstituting recombinant holophosphatases have generated two models, one proposing that substrate recruitment requires the addition of actin, whilst the second proposes that this function is encoded by R15s. The biological relevance of actin in substrate recruitment has not been evaluated. Here we generated a series of truncation mutants and tested their properties in mammalian cells. We show that substrate recruitment is encoded by an evolutionary conserved region in R15s, R15A^325-554^ and R15B^340-639^. Actin does not bind these regions establishing that it is not required for substrate recruitment. Activity assays in cells showed that R15A^325-674^ and R15B^340-713^, encompassing the substrate-binding region and the PP1 binding-region, exhibit wild-type activity. This study identifies essential regions of R15s and demonstrates they function as substrate receptors. This work will guide the design of future structural studies with biological significance.

## Introduction

Phosphorylation of the alpha subunit of the heterotrimeric translation initiation factor 2 (eIF2α) on serine 51 (1) is an evolutionarily conserved defence mechanism essential for cell survival and organismal fitness (2,3). It results in a transient reduction of bulk translation activity concomitant with a selective increase in translation of a subset of transcripts (1,4). In mammals, the phosphorylation levels of eIF2α are tuned to cellular needs through the antagonistic actions of four different eIF2α kinases, PKR, HRI, PERK and GCN2, and two eIF2α phosphatases (1). The latter are composed of the catalytic subunit PP1 bound to one of two related non-catalytic subunits PPP1R15A (5,6) or PPP1R15B (7). Because PP1 controls a large number of protein dephosphorylation events, it was initially proposed to be non-selective (8,9). However, whilst PP1 dephosphorylates many substrates *in vitro* (8,10), it is not free in cells, but bound to one or more proteins (9,11). Non-catalytic subunits of PP1 phosphatases are dissimilar but they bind PP1 using conserved short linear motifs, the most common of which is RVxF, that docks to a surface of PP1 opposite the catalytic site (12).

Previous studies have revealed the importance of various regions of R15A using cellular readouts. The function of human R15A was elucidated on the basis of its sequence similarity to the protein ICP34.5 from herpes simplex virus, which blocks the host response to infection by recruiting PP1 to dephosphorylate eIF2α (13). Further analyses led to the discovery that PP1 binding is through the carboxy-terminal region of ICP34.5, which harbours an RVxF (14). The carboxy-terminal region of ICP34.5 is essential for its function and can be replaced by the homologous region of R15A (14). A genetic screen in mammals later identified a fragment of R15A on the basis of its ability to repress expression of *Chop*, a target of the Integrated Stress Response (ISR) (5). This fragment, called A1, consists of hamster R15A^292-590^ (homologous to human R15A^310-674^) (5). Structure-function analyses revealed that the first ∼ 200 amino acids of R15A are not required for the ISR-repressing activity of the overexpressed protein but important for its localization to the ER (5,15,16). In contrast, the carboxy-terminal region is essential for function (5,15). Expression of the carboxy-terminal 70 amino acids was deemed to be active in repressing ISR target genes, although the activity of this small fragment was markedly decreased relative to hamster R15A^292-590^ or full-length mouse protein (5). Similar observations were made following overexpression of human R15 fragments in yeast expressing human eIF2α and PP1: full-length and R15A^420-674^ were equally active in rescuing from the toxicity resulting from overexpression of an eIF2α kinase (17). In the same system, R15A^513-674^ had residual activity (17). Importantly, active A1 is homologous to full-length ICP34.5 (13). Thus, these findings suggest that the first ∼ 200 amino-acids of R15A are dispensable for the function of the overexpressed protein.

We previously reconstituted human eIF2α phosphatases with recombinant PP1 and large fragments of recombinant R15A and R15B homologous to ICP34.5 (18). These complexes dephosphorylate eIF2α, but not other substrates, establishing that they recapitulated the function and selectivity of the native holoenzymes (18). In this minimal system, composed of R15, PP1 and the amino-terminal fragment of eIF2α, R15s recruit the substrate, providing substrate selectivity to the holoenzyme (18). The carboxy-terminal part of R15, with the RVxF, recruits PP1 but the resulting complex is not selective for eIF2α (18), in agreement with previous studies (15,19,20). The eIF2α binding region mapped to the middle of R15s and is essential for selectivity of the reconstituted enzymes (18). Another study also reported binding of eIF2α to the middle region of a recombinant R15A (19). According to these findings, a selective holophosphatase is a split enzyme that requires the assembly of two components, one harbouring substrate binding and the other, providing the catalytic function, both being essential (10).

In an alternative model, G-actin was proposed to provide substrate specificity to the otherwise unselective recombinant eIF2α phosphatases reconstituted with PP1 and the carboxy-terminal 70 amino acid region of R15s (20). This model stems from the observation that, overexpression of this region is sufficient to decrease expression of *CHOP*, an ISR target (5). However, its activity is marginal compared to longer fragments (5). Moreover, *in vitro*, this fragment does not confer substrate selectivity to the enzyme, which has similar properties to PP1 alone and dephosphorylates various substrates (20,21). This prompted the search for a selectivity factor and left two open questions: what is the function of R15s and why humans have 674 and 713 amino acid long R15A and R15B proteins. Actin co-precipitates with overexpressed GFP-tagged R15A (22). A crystal structure of a recombinant complex comprising of PP1, R15B^630-701^, and G-actin was reported (20). Actin was proposed to confer substrate selectivity in this *in vitro* system (20). However, the biological relevance of actin binding for substrate recruitment has not been examined so far.

The two current models for substrate recognition by eIF2α phosphatases agree that a selectivity factor is required in addition to the ∼70 amino-acid PP1-recruiting region. However, one model proposes that selective substrate recruitment requires longer fragments of R15s (18), and the other is dependent on actin (20). It is important to resolve these discrepancies and elucidate the molecular requirements for substrate recruitment by eIF2α phosphatases not only because these enzymes are central controllers of cellular fitness and drug targets (18,23–25), but also because such an advance could provide principles for elucidating substrate recognition of the many uncharacterised PP1 holoenzymes. The two current models for substrate recognition by eIF2α phosphatases have been generated using recombinant proteins. Because R15s are predicted to be intrinsically disordered, recombinant proteins may not necessarily have biologically relevant folds and properties. To circumvent this issue, we investigated the substrate binding requirements of R15s in cell-based assays and identified that their middle regions are responsible for substrate recruitment in absence of actin.

## Results

To identify functional regions in R15s, we generated a series of truncation mutants that were FLAG-tagged on their amino-termini. They were based on homology with the shorter viral protein ICP34.5, as well as clone A1 (Figure 1). Overexpressed R15A derivatives were immunoprecipitated with anti-FLAG antibodies (Figure 2a). PP1 was found to interact with the R15A derivatives that encompassed residues 554-674 (Figure 2a). eIF2α was enriched in the R15A^325-554^ and R15A^325-674^ pulldowns as well as in the pulldown of the full-length R15A^1-674^, albeit to a lesser extent (Figure 2a). We found no enrichment of eIF2α upon immunoprecipitation of R15A^1-325^ or R15A^554-674^ (Figure 2a). This reveals that R15A^325-554^ encodes the substrate recruitment region.

**Figure 1:**
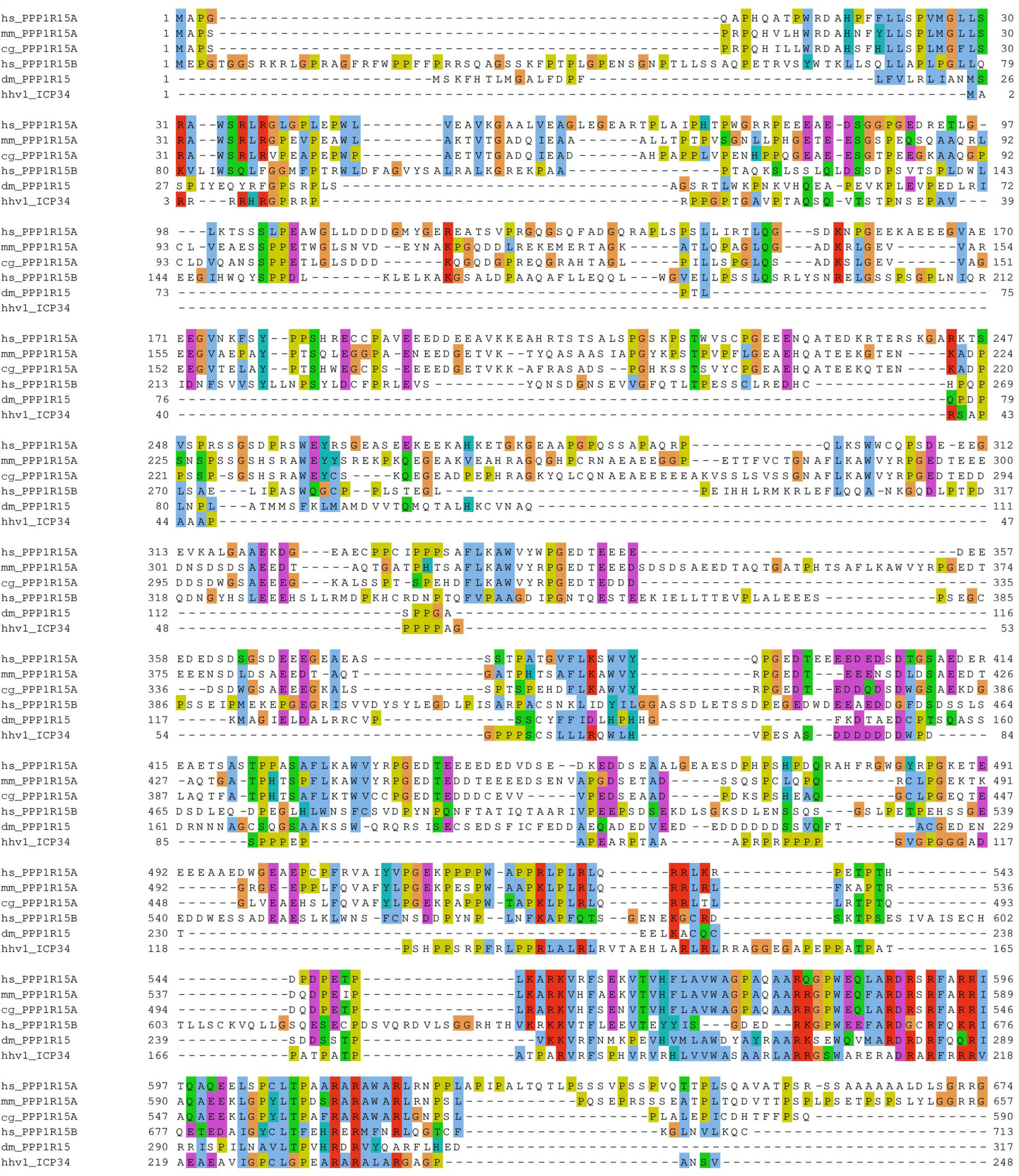
Multiple sequence alignment of various R15s. The sequences of R15A from H. sapiens (O75807), M. Musculus (P17564), C. griseus (A0A3L7HFU6), R15B from H. sapiens (Q5SWA1), R15 from, D. melanogaster (Q9W1E4) and ICP34 from Human herpesvirus 1 (P36313) were aligned using Muscle (33) and coloured according to CLUSTAL colouring scheme. The produced alignment was manually refined using Jalview (34).

**Figure 2:**
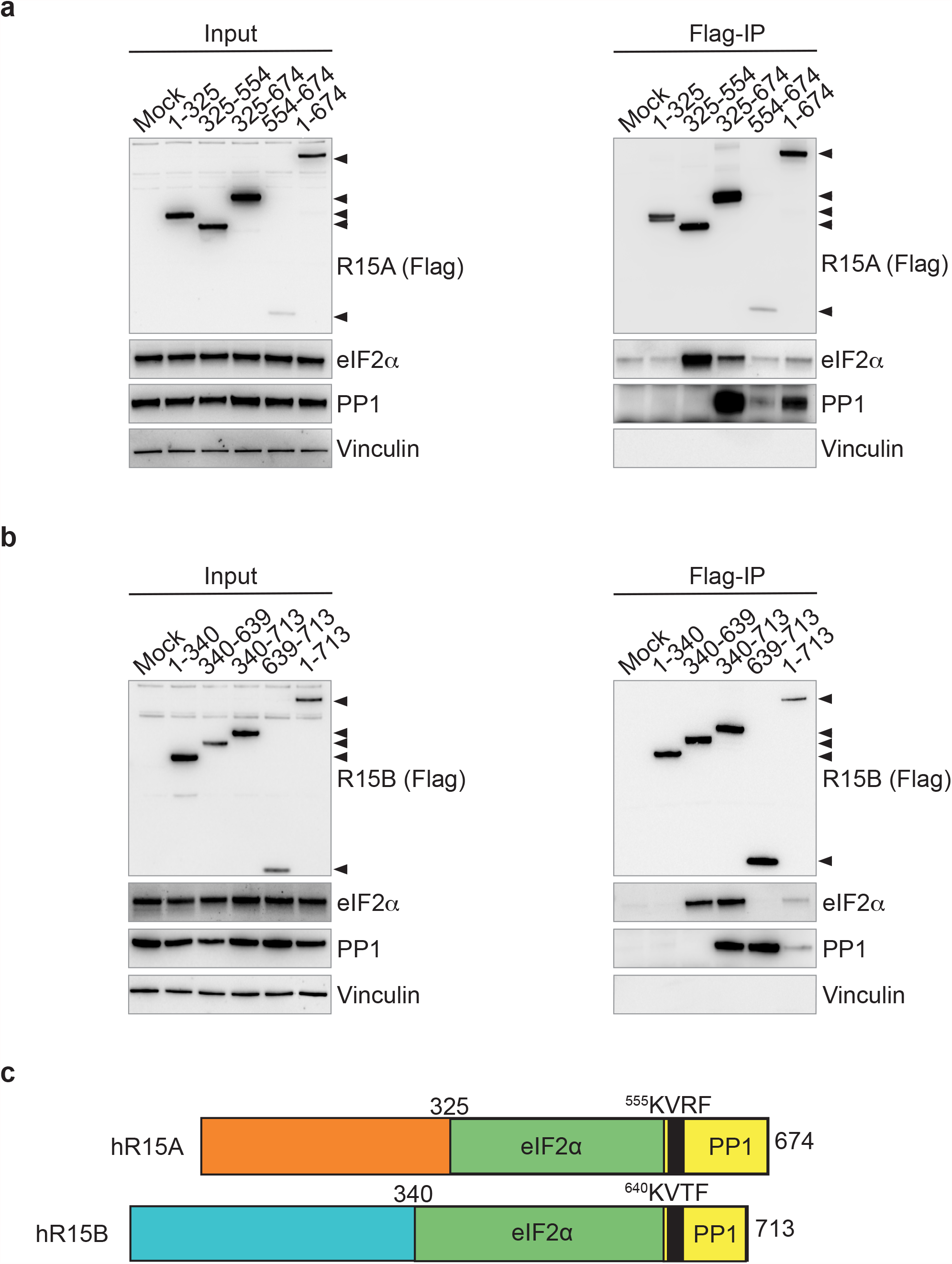
eIF2α and PP1 bind to different regions of R15s. (a, b) R15 constructs were transfected into HEK 293T cells (Input) and immunoprecipitated using anti-FLAG M2 magnetic beads (Flag-IP). Samples were eluted from the beads by boiling in LDS and eluates were separated on a 4-12% Bis Tris Plus gel. Proteins were detected by immunoblotting with FLAG, total eIF2α, PP1 and vinculin antibodies. (c) A cartoon summary of the modular binding of eIF2α and PP1 to both R15A and R15B. Representative results of at least 3 experiments are shown.

We next generated homologous truncation mutants of R15B (Figure 2b). R15B derivatives containing its carboxy-terminal region captured PP1 when immunoprecipitated (Figure 2b). In contrast, eIF2α was captured by R15B fragments (R15B^340-639^ and R15B^340-713^) that contained its middle region (Figure 2b). R15B^639-713^ co-precipitated PP1 but not eIF2α (Figure 2b). Conversely, R15B^340-639^ was bound to large amounts of eIF2α in absence of PP1 (Figure 2b). Full-length R15B captured both eIF2α and PP1 (Figure 2b). These experiments reveal that the binding of eIF2 and PP1 on R15B is not overlapping: R15B^340-639^ binds eIF2α whilst the carboxy-terminal region R15B^639-713^ binds PP1 (Figure 2b). Thus, R15s are modular proteins with their carboxy-termini binding PP1, whilst the substrate recruitment region is encoded by a distinct region in the middle of the proteins (Figure 2c).

Next we examined if substrate recruitment by eIF2α phosphatases depends on actin in cells. We found that actin was significantly enriched following immunoprecipitation of R15A^554-674^ (Figure 3a), in agreement with a previous report (22). As shown in Figure 2, this fragment did not capture eIF2α (Figure 3a). Conversely, the fragments that recruited the substrate did not pull-down actin (Figure 3a). This reveals that the actin-binding and the substrate-binding regions do not overlap. Full-length R15A did not capture actin (Figure 3a). Similar observations were made with R15B fragments. Actin was significantly captured following immunoprecipitation of R15B^639-713^ (Figure 3b) but the middle region of R15B (R15B^340-639^) captured eIF2α, in absence of actin (Figure 3b). These findings establish that that actin is not required for substrate recruitment.

**Figure 3:**
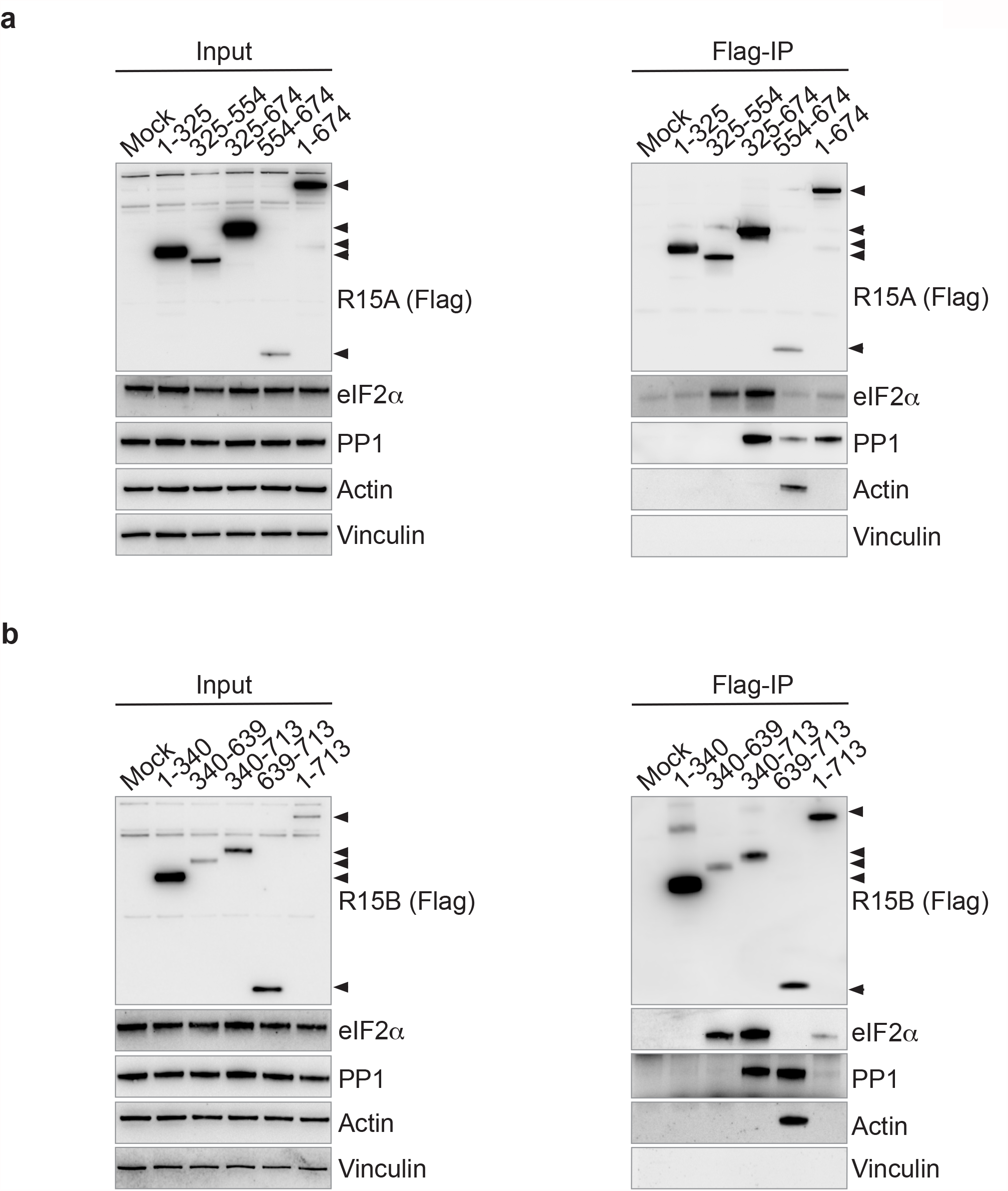
Actin binding is not required for substrate recruitment by R15A or R15B. (a, b) R15 constructs were transfected into HEK 293T cells (Input) and immunoprecipitated using anti-FLAG M2 magnetic beads (Flag-IP). Samples were eluted by boiling in LDS and eluates were separated on a 4-12% Bis Tris Plus gel. Proteins were detected by immunoblotting with FLAG, total eIF2α, PP1, Actin and vinculin antibodies. Representative results of at least 3 experiments are shown.

We next tested the ability of R15 fragments to dephosphorylate eIF2α in cells exposed to nutrient stress (1). Overexpression of full-length R15A^1-674^ led to a robust decrease in the levels of phosphorylated eIF2α (Figure 4a). Overexpression of R15A^325-674^ also led to nearly complete dephosphorylation of eIF2α (Figure 4a). Similarly, full-length R15B^1-713^ as well as R15B^340-713^ were active (Figure 4b). The amino-terminal regions of R15s were inactive, as well as R15A^325-554^ and R15B^340-639^ (Figure 4a and 4b). These results are in agreement with the properties of the recombinant holoenzymes (18). Intriguingly, overexpression of the carboxy-terminal fragment of R15A or R15B slightly but reproducibly decreased eIF2α phosphorylation (Figure 4a-b, 4c and Supplementary figure S1). Because these fragments bind actin as well as PP1 but not the substrate, we concluded that this activity is probably indirect. Indeed, these fragments are inactive *in vitro* (18,20).

**Figure 4:**
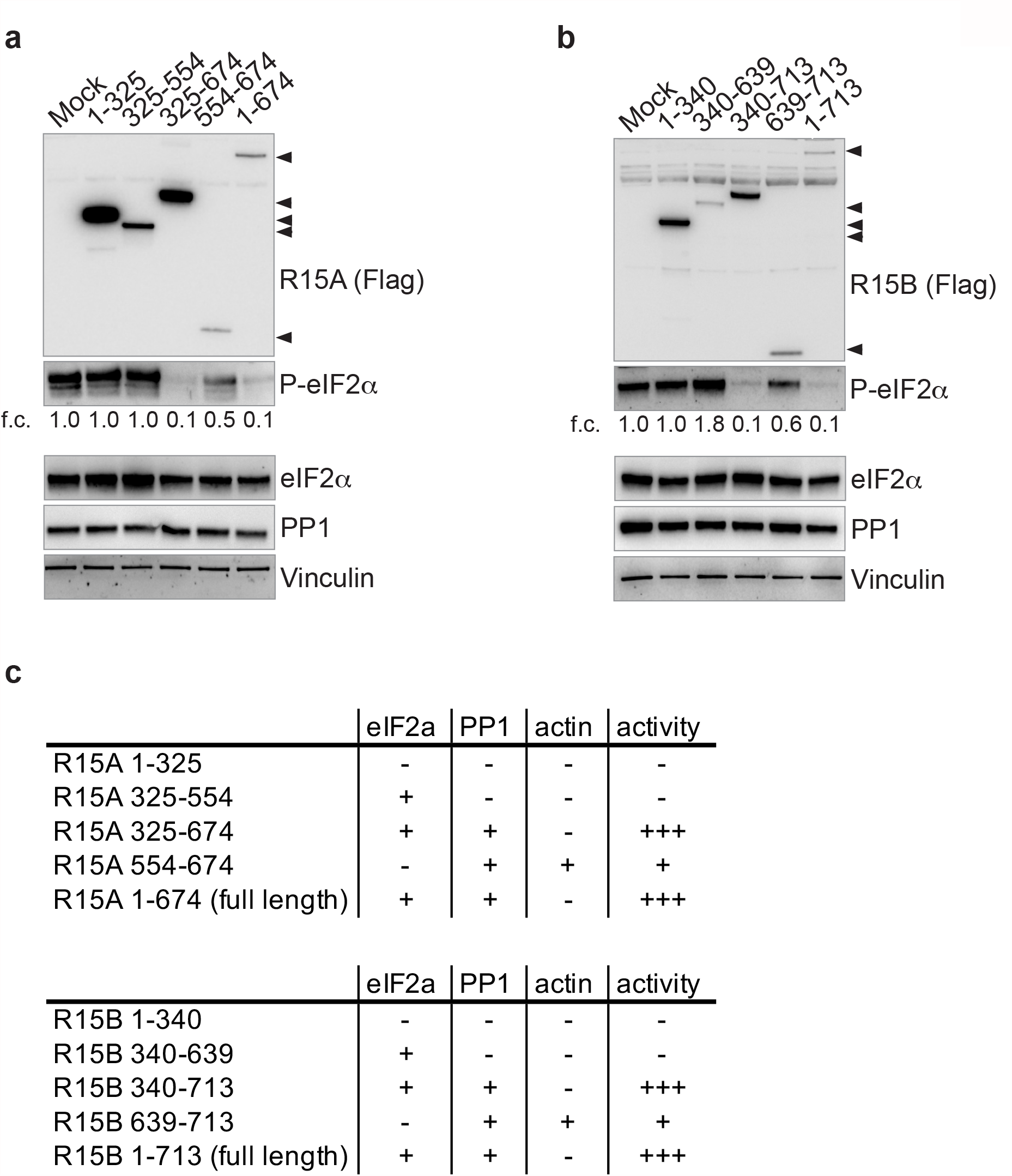
Activity of the various R15 constructs dephosphorylating P-eIF2α. (a, b) R15 constructs were transfected into HEK 293T cells, such that cells were overconfluent before lysis, providing high basal eIF2α phosphorylation to asses R15 construct activity. Proteins were detected by immunoblotting with FLAG, P-eIF2α, total eIF2α, PP1 and vinculin antibodies. Representative results of at least 3 experiments are shown. f.c.: Fold changes relative to mock transfection. (c) Table summarising the properties of R15 truncations.

To enable direct comparison with previously published fragments, we generated R15A^533-624^ used in (21), the hamster A1 fragment (5) as well as R15A^325-636^ (18) (Supplementary figure S1a) and compared their properties with the variants generated in this study. Transfection of R15A^533-624^ decreased levels of phosphorylated eIF2α in human cells to a small extend, similar to R15A^554-674^ fragment (Supplementary Figure S1b). In contrast, overexpressed A1 and R15A^325-636^ decreased eIF2α phosphorylation to background levels (Supplementary figure S1a), similar to R15A^325-674^ and R15B^340-713^ as well as full-length R15s. A1, R15A^325-636^, R15A^325-674^ and R15B^340-713^ bound eIF2 as well as PP1, unlike R15A^533-624^, which bound PP1 but not eIF2α (Supplementary figure S1a). These results reveal that full activity of all fragments tested correlates with substrate and PP1 binding capacity. This work defines R15A^325-674^ and R15B^340-713^ as active fragments, which bind both eIF2α and PP1 (Figure 4d).

## Discussion

We generated a series of truncation variants of R15A and R15B to study their properties in mammalian cells. We find that the amino-terminal ∼ 300 amino acids are not required for recruitment of eIF2α, PP1 or eIF2α dephosphorylation. We show that R15A^325-554^ and R15B^340-639^ are necessary and sufficient for the recruitment of eIF2α. Substrate recruitment does not require actin. These findings are compatible with prior studies of R15A in mammalian cells (15,19).

Whilst studies in cells provide a coherent body of information on the function of various regions of R15s, studies conducted *in vitro* with recombinant proteins have generated two models. In one model, substrate selectivity is provided by actin (20). In the second model, substrate selectivity is provided by the middle region of R15 (18). Here we compared the properties of these protein fragments in cells. As shown before (5,6,15,19), we observed here that the carboxy-terminal region of the proteins is necessary and sufficient to bind PP1. We show that substrate binding required the middle region of R15s. This finding is in agreement with an earlier study of R15A in cells (19) and an *in vitro* study (18). This region is evolutionarily conserved and present in ICP34.5 and clone A1.

We also assessed here the relevance of actin for substrate recruitment in cells. Whilst we validate the observation that the carboxy-terminal region of R15 is a strong actin binder (20,22), this property is restricted to short carboxy-terminal fragments of R15s. Actin could not be detected in pulldowns carried out with full-length proteins. Moreover, R15A^325-554^ and R15B^340-639^ capture eIF2α but not actin, establishing that substrate recruitment does not require actin.

Testing the activity of the various R15 truncation derivatives towards dephosphorylation of eIF2α yielded a coherent dataset. To be active, R15 fragments need to contain the middle region of the protein binding the substrate and the carboxy-terminal PP1 binding region (Figure 4c). These active fragments are homologous to clone A1 identified in an ISR suppressor screen (5). The amino-terminal region is not required for activity in cells (Figure 4) and *in vitro* (18) but important for subcellular localization (5,15,16). When expressed alone, the carboxy-terminal 70 amino acids of R15 bind PP1 and are marginally active. However, these fragments do not bind eIF2α, unlike the full-length protein.

The crystal structure of R15B^630-701^ bound to PP1 has revealed that R15B binds PP1 via its RVxF domain (20). The same group has deposited a follow-up crystallographic and cryoEM study where an amino-terminal fragment of eIF2α was added, as well Dnase1, to stabilize G-actin (26). In these studies, the biological requirement for actin in substrate recruitment was not evaluated. The findings reported here do not address the observation that various drugs altering actin dynamic induce eIF2α phosphorylation (22). Perturbation of actin polymerization has pleiotropic effects and is expected to induce stress responses, such as the ISR. However, the mechanism by which perturbation of actin polymerization impacts on eIF2α signalling is most likely indirect. We find that substrate recruitment does not require actin but is encoded by a region of R15s, that was missing from the fragments used in the structural work (20,26). We also show here that actin binding is a neomorphic property of the carboxy-terminal regions of R15s. Unlike the full-length proteins, which localize to the endoplasmic reticulum and does not bind actin, the small (< 100 aa) carboxy terminal fragments of R15s can freely diffuse into the nucleus (15), providing a possible explanation for their ability to bind G-actin.

The catalytic mechanism of PP1 dephosphorylation was elucidated more than two decades ago (27,28). There are only a couple of examples showing how non-catalytic subunits of phosphatases can directly support substrate recruitment (29,30). Structural studies with intrinsically disordered proteins such as R15s (19) are notoriously difficult. Deletion studies such as the one conducted here can be carried out with any non-catalytic subunits of PP1 to identify substrate-binding regions.

## Materials & Methods

### Cloning of R15 variants

All R15 derivatives were cloned from R15A/B human cDNA into the mammalian expression vector pXJ41 (31), which contains a single amino termini FLAG tag, using the Gibson Assembly method (32). Clone A1 was cloned from haA1.pBABEpu. Primer sequences used are available in Supplementary table 1

All R15 constructs cloned into the PXJ41 mammalian expression vector, harbouring a single amino termini FLAG tag.

| **R15A Constructs** | **Species** | **Boundaries based upon:** |
| --- | --- | --- |
| 1-325 | Human | Conservation and Carrara et al., 2017 |
| 325-554 | Human | Conservation and Carrara et al., 2017 |
| 325-636 | Human | Carrara et al., 2017 |
| 325-674 | Human | Conservation and Carrara et al., 2017 |
| 554-674 | Human | Conservation and Carrara et al., 2017 |
| 1-674 (Full-length) | Human | Conservation and Carrara et al., 2017 |
| 533-624 | Human | Chen et al., 2015 and Crespillo-Casado et al., 2017 |
| A1 292-590 | Hamster | Novoa et al., 2001<br>Corresponding to 311-658 Human R15A |

| **R15B Constructs** | **Species** | **Boundaries based upon:** |
| --- | --- | --- |
| 1-340 | Human | Conservation and Carrara et al., 2017 |
| 340-639 | Human | Conservation and Carrara et al., 2017 |
| 340-713 | Human | Conservation and Carrara et al., 2017 |
| 639-713 | Human | Conservation and Carrara et al., 2017 |
| 1-713 (Full-length) | Human | Conservation and Carrara et al., 2017 |

### Cell Culture

Human embryonic kidney 293T cells (HEK 293T) were grown in a humidified incubator with 5% CO_2_ at 37°C. Cells were maintained in Dulbecco’s Modified Eagle’s Media (DMEM, Sigma, D5796) supplemented with 100 U/mL penicillin and 100 μg/mL streptomycin (Gibco, 15140122), 2 mM L-glutamine (Gibco, 25030) and 10% fetal bovine serum (FBS, Gibco 10270).

### Co-immunoprecipitation assay

#### Transient PEI transfection

HEK 293T cells were seeded at 0.8×10^6^ cells per 10 cm^2^ dish and grown for 24 hours prior to transfection. 4 µg of the indicated R15 constructs were added to 800 µl of Opti-MEM media (Gibco, 11058) and mixed. Subsequently, 12 µl PEI transfection reagent (Polysciences, 24765) was added, mixed and incubated 20 minutes at room temperature (RT). The transfection mixture was then added to the 10cm^2^ cell culture dish drop wise and mixed gently. Cells were incubated for an additional 24 hours.

#### Cell lysates

Cells were gently washed with 5 ml ice-cold PBS before collected and pelleted at 300 RCF for 5 minutes. The cell pellet was lysed in 800 µl of lysis buffer (50 mM Tris-HCl pH 7.4, 10 mM sodium chloride, 100 mM potassium chloride, 0.1 µM calcium chloride, 0.5 mM magnesium chloride, 0.5 mM TCEP, EDTA-free complete protease tablet (Roche, 04639159001)). The lysates were sonicated on ice for 3x 3 seconds using a Microson ultrasonic cell disrupter XL (Misonix), with an output power of 0.06 Watts (RMS Watts). The lysates were clarified by microcentrifugation at 4 °C for 12 minutes at 16000 RCF. Supernatants were transferred to fresh tubes.

#### FLAG Immunoprecipitation

For each condition, 10 µl anti-FLAG M2 magnetic beads (Sigma-Aldrich, M8823-1ML), pre-equilibrated in lysis buffer (see above), were added to 700 µl of lysates and incubated overnight at 4 °C on a rotating wheel. Samples were washed three times with lysis buffer and proteins were eluted upon addition of 50 µl of 1X BOLT LDS (Novex #B007) with 100 mM DTT and boiling at 95 °C for 10 minutes. 10 µl of the immunoprecipitated samples as well as 10 µl of lysates (input) were analysed by immunoblots.

### Immunoblotting

Proteins were separated on Bolt 4-12% Bis-Tris Plus gel (Invitrogen, #NW04120BOX) in 1X MES running buffer. 2 µL of Protein Precision Plus Dual Colour Standards (#161-0374) was loaded on each gel. Gels were run at 120 V for 70 minutes and transferred onto a nitrocellulose membrane (Bio-Rad, 1704159) using a Trans-Blot Turbo System (BioRad). All membranes were stained using Ponceau S (Sigma, P7170) solution for 3 minutes to assess transfer quality and equal loading. Membranes were then blocked for 1 hour at RT using 5% milk in TBS with 0.025% Tween 20 (Sigma, P1379) (TBS-T) with shaking. Membranes were rinsed 3 times with TBS-T and incubated with the relevant primary antibody diluted in 5% BSA in TBS-T overnight at 4 °C, while shaking. Membranes were then washed 3 times with TBS-T before incubating with the relevant secondary antibody in TBS-T with 5% milk for 1 hour at RT while shaking. Following being washed 3 times with TBS-T and once with TBS, Amersham ECL Prime detection reagent kit (GE Healthcare Life Sciences, RPN2232) was used to detect chemiluminescence with ChemiDoc Touch Imaging System (Bio-Rad).

### Activity Assay

HEK 293T cells were seeded at 1.5×10^6^ to provide overconfluent cells at the time of harvesting. The day after seeding, cells were transfected and lysed as described above.

### Antibodies

| Protein | Antibody | Concentration used | Species |
| --- | --- | --- | --- |
| PP1 | Sc-7482 | 1:1000 | Mouse |
| Total eIF2α | ab26197 | 1:1500 | Rabbit |
| P-eIF2α | 44-728G | 1:1000 | Rabbit |
| Vinculin | 4650S | 1:5000 | Rabbit |
| FLAG | F7425 | 1:1000 | Rabbit |
| Actin | 20536-1-AP | 1:10000 | Rabbit |
| Anti-Rabbit IgG (H+L), HRP Conjugate | W4011 | 1:10000 | Goat |
| Anti-Mouse IgG (H+L), HRP Conjugate | W4021 | 1:10000 | Goat |

## Acknowledgements

We are grateful to Marta Carrara, Milena Krach, Marios Koliopoulos, Max Dalglish for pilot studies revealing some of the findings reported here using various assays as well as Annika Weber for the cellular activity assay. haA1.pBABEpu was a gift from David Ron (Addgene plasmid # 21813; http://n2t.net/addgene:21813; RRID:Addgene_21813). We thank Maria Szaruga for help with the manuscript and Michel Goedert for comments on the manuscript.

## Author’s contribution

AB and GH designed the study, GH conducted the experiments, analysed the data and prepared figures. AA carried out sequence alignments. AB wrote the manuscript.

## Figure legends

**Supplementary Figure 1.**
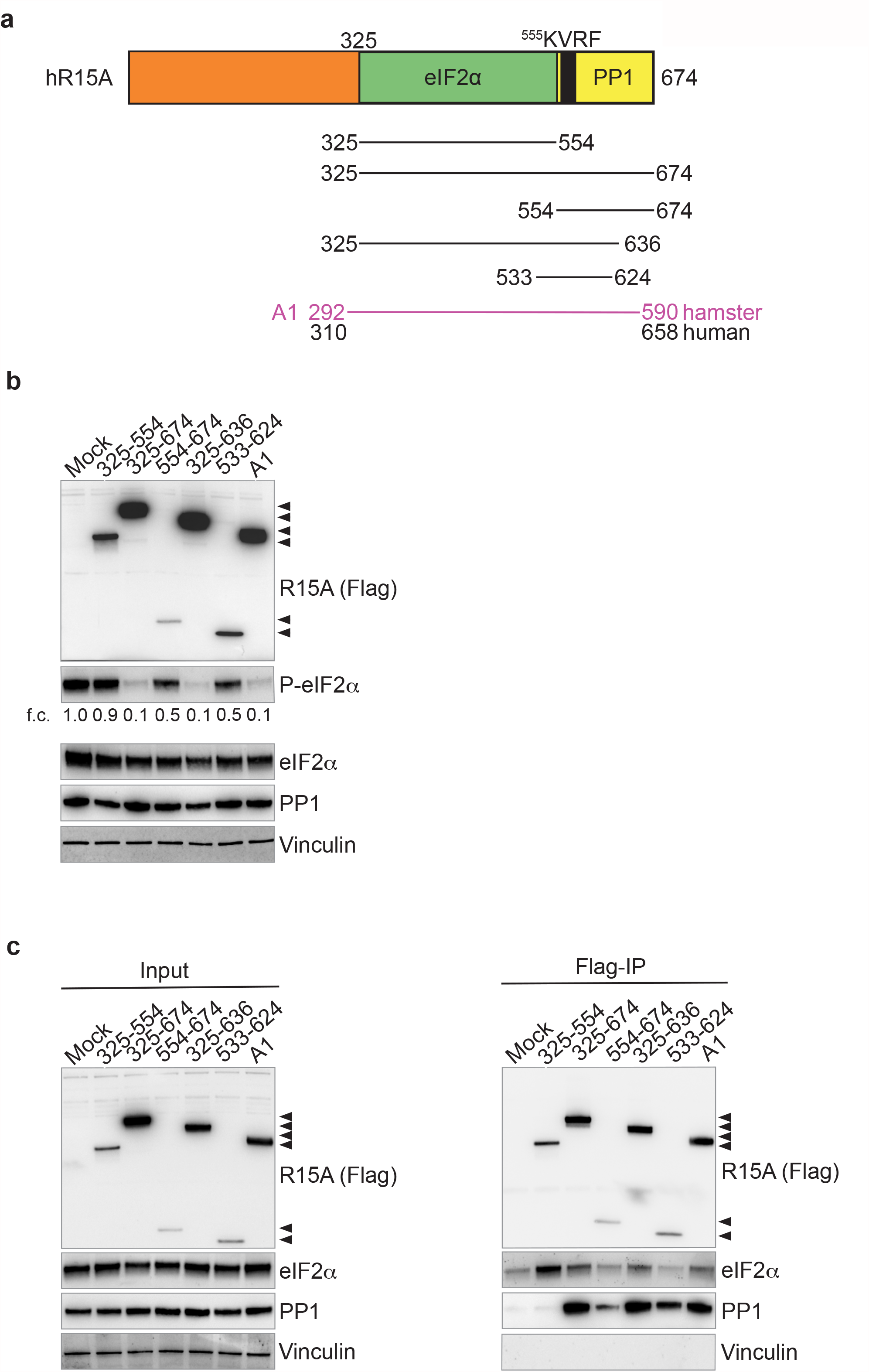
Properties of previously published R15s. (a) Cartoons of the proteins. The amino acid position of hamster clone A1 are indicated in magenta and the corresponding positions of the human proteins are indicated in black. All other proteins are human. (b) Activity of transfected R15 derivatives assessed by decreased levels of P-eIF2α in cells cultured with high basal level of P-eIF2α. f.c.: Fold changes relative to mock transfection. (c) R15 constructs were transfected into HEK 293T cells (Input) and immunoprecipitated using anti-FLAG M2 magnetic beads (FLAG-IP). Immunoprecipitated complexes were eluted by boiling in LDS and separated on a 4-12% Bis Tris Plus gel. Proteins were detected by immunoblotting with FLAG, total eIF2α, PP1, and vinculin antibodies. Representative results of at least 3 experiments are shown.

**Supplementary Table 1.**
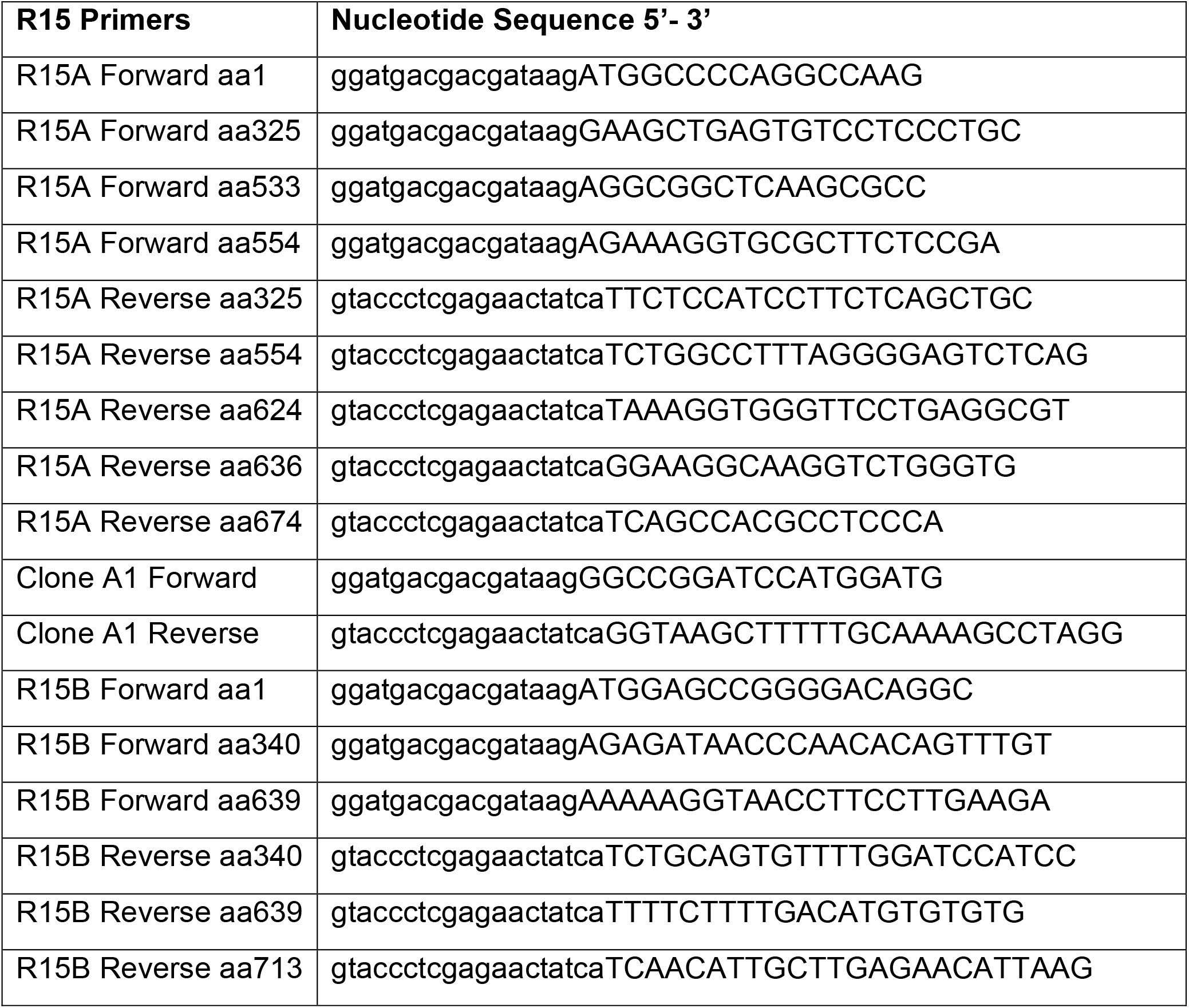
Primers used for cloning R15 constructs. Nucleotides in lower case represent the overhangs used for the insertion into PXJ41, via the Gibson Assembly procedure.

## References

1. Wek RC. Role of eIF2α Kinases in Translational Control and Adaptation to Cellular Stress. Csh Perspect Biol. 2018;10(7):a032870.

2. Luh LM, Bertolotti A. Potential benefit of manipulating protein quality control systems in neurodegenerative diseases. Curr Opin Neurobiol. 2020;61:125–32.

3. Costa-Mattioli M, Walter P. The integrated stress response: From mechanism to disease. Science. 2020;368(6489):eaat5314.

4. Schneider K, Nelson GM, Watson JL, Morf J, Dalglish M, Luh LM, et al. Protein Stability Buffers the Cost of Translation Attenuation following eIF2α Phosphorylation. Cell Reports. 2020;32(11):108154.

5. Novoa I, Zeng H, Harding HP, Ron D. Feedback Inhibition of the Unfolded Protein Response by GADD34-Mediated Dephosphorylation of eIF2α. J Cell Biology. 2001;153(5):1011–22.

6. Connor JH, Weiser DC, Li S, Hallenbeck JM, Shenolikar S. Growth Arrest and DNA Damage-Inducible Protein GADD34 Assembles a Novel Signaling Complex Containing Protein Phosphatase 1 and Inhibitor 1. Mol Cell Biol. 2001;21(20):6841– 50.

7. Jousse C, Oyadomari S, Novoa I, Lu P, Zhang Y, Harding HP, et al. Inhibition of a constitutive translation initiation factor 2α phosphatase, CReP, promotes survival of stressed cells. J Cell Biology. 2003;163(4):767–75.

8. Virshup DM, Shenolikar S. From Promiscuity to Precision: Protein Phosphatases Get a Makeover. Mol Cell. 2009;33(5):537–45.

9. Roy J, Cyert MS. Cracking the Phosphatase Code: Docking Interactions Determine Substrate Specificity. Sci Signal. 2009;2(100):re9–re9.

10. Bertolotti A. The split protein phosphatase system. Biochem J. 2018;475(23):3707–23.

11. Hendrickx A, Beullens M, Ceulemans H, Abt TD, Eynde AV, Nicolaescu E, et al. Docking Motif-Guided Mapping of the Interactome of Protein Phosphatase-1. Chem Biol. 2009;16(4):365–71.

12. Egloff M, Johnson DF, Moorhead G, Cohen PTW, Cohen P, Barford D. Structural basis for the recognition of regulatory subunits by the catalytic subunit of protein phosphatase 1. Embo J. 1997;16(8):1876–87.

13. He B, Gross M, Roizman B. The γ134.5 protein of herpes simplex virus 1 complexes with protein phosphatase 1α to dephosphorylate the α subunit of the eukaryotic translation initiation factor 2 and preclude the shutoff of protein synthesis by double-stranded RNA-activated protein kinase. Proc National Acad Sci. 1997;94(3):843–8.

14. He B, Gross M, Roizman B. The γ134.5 Protein of Herpes Simplex Virus 1 Has the Structural and Functional Attributes of a Protein Phosphatase 1 Regulatory Subunit and Is Present in a High Molecular Weight Complex with the Enzyme in Infected Cells* . J Biol Chem. 1998;273(33):20737–43.

15. Brush MH, Weiser DC, Shenolikar S. Growth Arrest and DNA Damage-Inducible Protein GADD34 Targets Protein Phosphatase 1α to the Endoplasmic Reticulum and Promotes Dephosphorylation of the α Subunit of Eukaryotic Translation Initiation Factor 2. Mol Cell Biol. 2003;23(4):1292–303.

16. Kloft N, Neukirch C, Hoven G von, Bobkiewicz W, Weis S, Boller K, et al. A Subunit of Eukaryotic Translation Initiation Factor 2α-Phosphatase (CreP/PPP1R15B) Regulates Membrane Traffic* . J Biol Chem. 2012;287(42):35299–317.

17. Rojas M, Vasconcelos G, Dever TE. An eIF2α-binding motif in protein phosphatase 1 subunit GADD34 and its viral orthologs is required to promote dephosphorylation of eIF2α. Proc National Acad Sci. 2015;112(27):E3466–75.

18. Carrara M, Sigurdardottir A, Bertolotti A. Decoding the selectivity of eIF2α holophosphatases and PPP1R15A inhibitors. Nat Struct Mol Biol. 2017;24(9):708– 16.

19. Choy MS, Yusoff P, Lee IC, Newton JC, Goh CW, Page R, et al. Structural and Functional Analysis of the GADD34:PP1 eIF2α Phosphatase. Cell Reports. 2015;11(12):1885–91.

20. Chen R, Rato C, Yan Y, Crespillo-Casado A, Clarke HJ, Harding HP, et al. G-actin provides substrate-specificity to eukaryotic initiation factor 2α holophosphatases. Elife. 2015;4:e04871.

21. Crespillo-Casado A, Chambers JE, Fischer PM, Marciniak SJ, Ron D. PPP1R15A-mediated dephosphorylation of eIF2α is unaffected by Sephin1 or Guanabenz. Elife. 2017;6:e26109.

22. Chambers JE, Dalton LE, Clarke HJ, Malzer E, Dominicus CS, Patel V, et al. Actin dynamics tune the integrated stress response by regulating eukaryotic initiation factor 2α dephosphorylation. Elife. 2015;4:e04872.

23. Tsaytler P, Harding HP, Ron D, Bertolotti A. Selective Inhibition of a Regulatory Subunit of Protein Phosphatase 1 Restores Proteostasis. Science. 2011;332(6025):91–4.

24. Das I, Krzyzosiak A, Schneider K, Wrabetz L, D’Antonio M, Barry N, et al. Preventing proteostasis diseases by selective inhibition of a phosphatase regulatory subunit. Science. 2015;348(6231):239–42.

25. Krzyzosiak A, Sigurdardottir A, Luh L, Carrara M, Das I, Schneider K, et al. Target-Based Discovery of an Inhibitor of the Regulatory Phosphatase PPP1R15B. Cell. 2018;174(5):1216-1228.e19.

26. Yan Y, Harding HP, Ron D. Higher order phosphatase-substrate contacts terminate the Integrated Stress Response. Biorxiv. 2021;2021.06.18.449003.

27. Goldberg J, Huang H, Kwon Y, Greengard P, Nairn AC, Kuriyan J. Three-dimensional structure of the catalytic subunit of protein serine/threonine phosphatase-1. Nature. 1995;376(6543):745–53.

28. Egloff M-P, Cohen PTW, Reinemer P, Barford D. Crystal Structure of the Catalytic Subunit of Human Protein Phosphatase 1 and its Complex with Tungstate. J Mol Biol. 1995;254(5):942–59.

29. Terrak M, Kerff F, Langsetmo K, Tao T, Dominguez R. Structural basis of protein phosphatase 1 regulation. Nature. 2004;429(6993):780–4.

30. Fedoryshchak RO, Přechová M, Butler A, Lee R, O’Reilly N, Flynn HR, et al. Molecular basis for substrate specificity of the Phactr1/PP1 phosphatase holoenzyme. Elife. 2020;9:e61509.

31. Xiao JH, Davidson I, Matthes H, Garnier J-M, Chambon P. Cloning, expression, and transcriptional properties of the human enhancer factor TEF-1. Cell. 1991;65(4):551–68.

32. Gibson DG, Young L, Chuang R-Y, Venter JC, Hutchison CA, Smith HO. Enzymatic assembly of DNA molecules up to several hundred kilobases. Nat Methods. 2009;6(5):343–5.

33. Edgar RC. MUSCLE: multiple sequence alignment with high accuracy and high throughput. Nucleic Acids Res. 2004;32(5):1792–7.

34. Clamp M, Cuff J, Searle SM, Barton GJ. The Jalview Java alignment editor. Bioinformatics. 2004;20(3):426–7.

